# *Mycobacterium tuberculosis, Mycobacterium kansasii* and *Rhodococcus equi* induce macrophage necroptosis in the presence of a caspase inhibitor acting on a non-canonical target(s)

**DOI:** 10.1101/2025.09.02.673674

**Authors:** Rong Hu, Pei Li, Jie Han, Jiamin Huang, Guoyi Huang, Xiuju Jiang, Daniel Pfau, Guobao Li, Yi-Nan Gong, Carl F. Nathan, Li Zhang

## Abstract

Macrophages are the predominant cell type infected by *Mycobacterium tuberculosis* (Mtb) in tuberculosis (TB). Death of Mtb-infected macrophages promotes tissue pathology and releases Mtb to infect other cells, suggesting that inhibiting the death of Mtb-infected macrophages could be an adjunctive treatment of TB. Prospects for such an intervention depend on identifying the molecular pathways leading to cell death. We previously reported that the death of Mtb-infected mouse macrophages in vitro depends on type I interferon (IFN) and that the ensuing upregulation of *cis*-aconitate decarboxylase (ACOD1; IRG1) contributes to cell death by exacerbating Mtb-induced lysosomal membrane permeabilization. Here we report that death of Mtb-infected primary mouse macrophages in vitro became necroptotic and die faster in the presence of benzyloxycarbonyl-Val-Ala-Asp-fluoromethylketone (z-VAD) acting on a target other than caspase-8. Macrophages infected with *Mycobacterium kansasii* and *Rhodococcus equi* likewise underwent z-VAD-dependent necroptosis. In C57BL/6 mice, which are relatively TB-resistant, we saw no impact of MLKL deficiency on bacterial burden or pulmonary pathology. In contrast, in *Sp140*^−/−^ mice on the C57BL/6 background, which express high levels of type I IFN after Mtb infection and develop necrotic pulmonary lesions, MLKL-deficiency reduced bacterial burden and pathology after high-dose infection. This report illustrates that off-target action(s) of a caspase-8 inhibitor can switch the cell death pathway to necroptosis in macrophages infected with various Gram-positive pathogens. In turn, this opens the possibility that pathophysiologic circumstances may lead to inhibition of the same target(s) that z-VAD inhibited in our studies. That may be what allows MLKL to exacerbate tuberculosis in mice that are prone to formation of necrotic lesions.

**Author Summary:** Tuberculosis (TB), caused by the bacillus *Mycobacterium tuberculosis* (Mtb), led to about 1.3 million deaths in 2023. The rapid emergence of drug resistance and slow pace of new drug development prompt attention to adjunctive host-directed therapies, an approach that relies on a thorough and detailed understanding of host-Mtb interactions. Mtb infection of macrophages can kill them, promoting tissue pathology and releasing of Mtb to infect other cells. Here we find that a pan-caspase inhibitor z-VAD hastened the death of Mtb-infected macrophages and converted the mode of cell death to RIPK1-RIPK3-MLKL-dependent necroptosis. However, the functional target(s) of z-VAD-FMK and the activation mechanism of RIPK3 differ from the canonical pathways. The pan-caspase inhibitor also promoted rapid, necroptotic death of macrophages infected with *Mycobacterium kansasii* and *Rhodococcus equi*. We further found that MLKL-deficiency in *Sp140*^−/−^ mice on the C57BL/6 background resulted in less weight loss, lower Mtb burdens and mitigation of lung pathology after high-dose Mtb infection. Our findings suggest that necroptosis can contribute to TB pathogenesis in some circumstances. Further identification of the death pathway-switching mechanism could deepen our understanding of host-Mtb interactions and aid in the design of adjunctive host-directed therapies.

## Introduction

Like many pathogens, Mtb manipulates important host cellular processes, including cell death pathways, for its own propagation. Virulent Mtb inhibits apoptosis of macrophages(1, 2), the initial and major cellular niche for Mtb infection and replication. Nonetheless, macrophages that fail to clear Mtb eventually die, releasing bacilli that can infect other cells in the same host or be expelled in aerosols to infect others. In addition, release of cellular content contributes to tissue inflammation.

Necroptosis is a type of programmed necrosis that classically requires receptor interacting protein kinase 1 (RIPK1), RIPK3 and mixed lineage kinase domain-like (MLKL)(3–6). Necroptosis has been most thoroughly studied in response to TNFα. TNFα signaling usually triggers NF-κB and mitogen-activated protein (MAP) kinases pathways that promote cell survival. However, if the activity of cellular inhibitor of apoptosis proteins (cIAPs) is inhibited by the second mitochondria-derived activator of caspases (Smac) protein or small molecule mimetics of Smac (Smac mimetics), caspases are activated, leading to apoptosis. If caspases are also inhibited, the death mode shifts to necroptosis, provided that the downstream necroptosis machinery is intact. RIPK1 and RIPK3, interacting through their respective homotypic interaction motifs (RHIMs), join with MLKL to form the necrosome. The interaction between RIPK1 and RIPK3 drives the autophosphorylation of RIPK3, enabling its recruitment and phosphorylation of MLKL. Phosphorylated MLKL oligomerizes and forms a pore in phospholipid-rich cell membranes, resulting in cell death(3–6). Necroptosis can also ensue in response to TLR signaling or viral infection(7, 8). In the latter two cases, the RHIM domain-containing proteins TRIF and ZBP1 (aka DAI), respectively, mediate the interaction with RIPK3. It was recently reported that mitochondrial DNA is released into the cytosol upon necroptosis induction and activates the cGAS-STING pathway to promote type I IFN expression(9). In vivo, necroptosis has been found in disorders and states ranging from nerve injury to senescence of the mouse male reproductive system(10–13).

Whether Mtb infection results in necroptosis is inconclusive. In one study, macrophages were protected from Mtb-induced cell death in vitro by depletion of RIPK3 or MLKL, and the numbers of Mtb and neutrophils in the lungs were modestly reduced in *Ripk3* knockout mice compared to wild type mice after high-dose intravenous infection(14). However, in other studies, low-dose aerosol infection led to a similar course of disease in wild type, *Mlkl*-deficient and *Ripk3*-deficient C57BL/6 mice(15, 16). Tuberculosis necrotizing toxin (TNT) is secreted by Mtb into macrophages, where its nicotinamide adenine dinucleotide glycohydrolase activity led to necroptosis that was dependent on RIPK3 and MLKL but not on TNFα or RIPK1 (17). However, Mtb lacking TNT was not defective for the course of infection in mice(18). In another study, components of the necroptotic machinery were increased in macrophages after Mtb infection both in vivo and in vitro, due in part to the type I IFN signaling induced by the infection. However, neither chemical nor genetic perturbation of the necroptosis pathway reduced the death of Mtb-infected macrophages(15, 16, 19). *Mycobacterium marinum* infection of zebrafish or cells of a human macrophage cell line led to a form of necrosis dependent on TNFα and production of mitochondrial reactive oxygen species (ROS), without relying on RIPK1, RIPK3 or MLKL(20).

In the present study, as before(19), we continued to observe that Mtb infection alone did not elicit necroptosis in mouse macrophages in vitro. However, addition of z-VAD, a presumed pan-caspase inhibitor, led to rapid cell death by necroptosis in a manner dependent on type I IFN, TNFα, RIPK1, RIPK3 and MLKL, but not dependent on inhibition of caspases. The Gram-positive pathogens *M. kansasii* and *R. equi* also elicited macrophage necroptosis in the presence of z-VAD. Finally, we observed that MLKL contributes to TB pathology in high-dose infection of mice prone to formation of necrotic tuberculous lesions. The evidence highlights that necroptosis of macrophages infected with various pathogens can be elicited in vitro through pharmacologic action on a target(s) other than caspase-8 and can occur in vivo, depending on the genetic background of the host.

## Results

### Mtb infection in the presence of z-VAD leads to macrophage necroptosis

Bone marrow-derived macrophages (BMDMs) from C57BL/6 mice died within one day post infection (d.p.i.) with Mtb if they were also exposed to the pan-caspase inhibitor z-VAD, a time at which death was not yet evident in macrophages infected with Mtb alone (Figures 1A and 1B). However, the caspase-1 inhibitor Ac-YVAD-CMK (YVAD), the caspase-8 inhibitor z-IETD-FMK (IETD) and another pan-caspase inhibitor, Q-VD-OPH, had no such effect (Figures 1A, S1A and S1B). The RIPK1 inhibitor necrostatin 1 (Nec-1) and the RIPK3 inhibitor GSK’872 that potently inhibit RIPK3 kinase activity(21) reverted the death phenotype seen with z-VAD (Figure 1B) without themselves exerting a toxic effect (Figure S1C). Consistent with the cell death in this setting being necroptotic, Mtb infection combined with z-VAD treatment triggered time-dependent phosphorylation of RIPK1 and MLKL (Figure S1D). Nec-1 inhibited both phosphorylation events, while GSK’872 inhibited phosphorylation of MLKL but not RIPK1 (Figure 1C and S1D). Moreover, a loss-of-function mutation in RIPK3 and deficiency of MLKL each conferred resistance to z-VAD-dependent, Mtb-related necroptosis (Figures 1D, 1E and S1E), as well as to necroptosis in response to the classical necroptosis-promoting combination of TNFα, Smac-mimetic and z-VAD (hereafter called TSZ) without Mtb (Figures S1F and S1G). The combination of TNFα, Smac-mimetic and Q-VD-OPH (hereafter called TSQ) triggered slower necroptosis than TSZ, but GSK’872 blocked necroptosis in both cases (Figure S1H).

**Figure 1.**
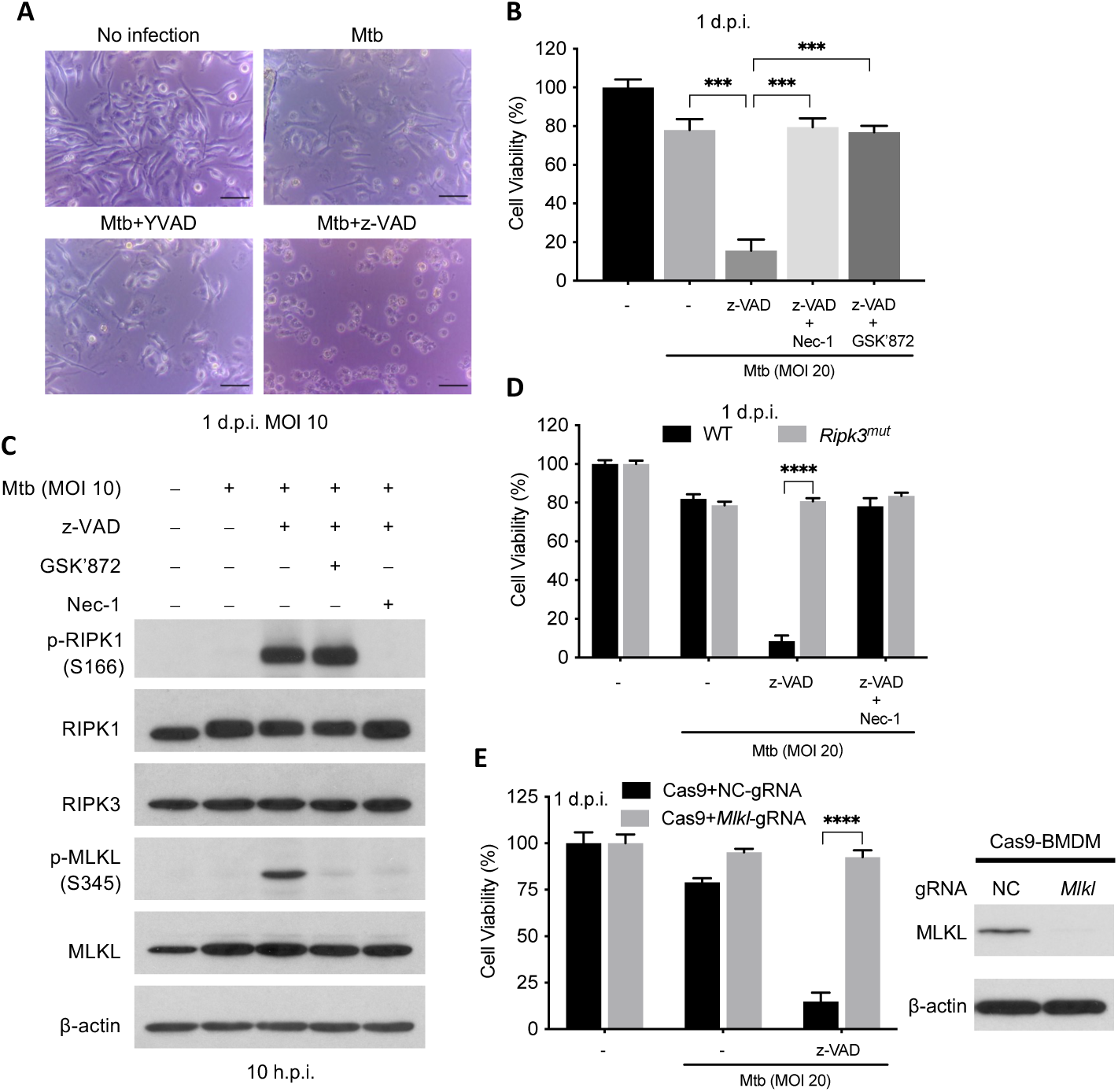
Mtb infection in the presence of z-VAD treatment induced RIPK1-RIPK3-MLKL-dependent necroptosis. BMDMs were pretreated with compounds at half of the indicated concentration for 2 h before infection of Mtb and then indicated concentration (20 μM except that GSK’872 is used at 3 μM) after washing Mtb away till harvest. (A) Photomicrographs of cells 1 d.p.i. Scale bar, 50 μm. (B) Cell viability measured by ATP assay 1 d.p.i. (C) Effects of RIPK1 and RIPK3 inhibitors on necroptotic activation of Mtb + z-VAD. Shown are immunoblots of cell lysates harvest at 10 h.p.i. (D and E) Effects of RIPK3 and MLKL deficiency on Mtb-related macrophage necroptosis. Cell viability was measured by ATP assay on 1 d.p.i. Data are means ± SD of three technical replicates (B, D and E) and representative of two experiments. ***, p < 0.001; ****, p< 0.0001 (two-tailed unpaired Student’s *t* test).

Even though Mtb + z-VAD robustly induced phosphorylation of mouse MLKL at Ser345, these stimuli did not induce detectable phosphorylation of mouse RIPK3 at Thr231 and Ser232(Figure S1I). In contrast, both sites were phosphorylated in response to TSZ (Figure S1I). Thus, Mtb-related, z-VAD-dependent necroptosis and TSZ-stimulated necrosis involve RIPK3 and its kinase activity but lead to its differential post-translational modification.

### Mtb-related, z-VAD-dependent necroptosis relies on TLR2-TNFα signaling as well as type I IFN signaling induced by infection

Classically, necroptosis involves TNFα signaling. Mtb-infected macrophages produce TNFα predominantly in response to a TLR2-MYD88-NF-κB pathway(22–24) and respond to it in autocrine or paracrine fashion through TNF receptor 1 (TNFR1). Consistent with this, we found that BMDMs deficient in TNFα or TNFR1 were resistant to Mtb-related, z-VAD-dependent necroptosis (Figure 2A, S2A1 and S2B). Likewise, genetic loss of TLR2, the major TLR for recognizing Mtb, or MYD88, a key adaptor downstream of TLR2, abolished the production of TNFα after Mtb infection (Figure 2B and S2C) and prevented Mtb-related necroptosis (Figure 2A, S2D and S2E). In contrast, TLR4, 5, 7 and 9 and adaptor proteins TRIF and Sarm1 (a.k.a. Myd88-5) were each dispensable for necroptosis (Figure 2A, S2D-H) and TRIF was dispensable for TNFα production after Mtb infection (Figure 2B).

**Figure 2.**
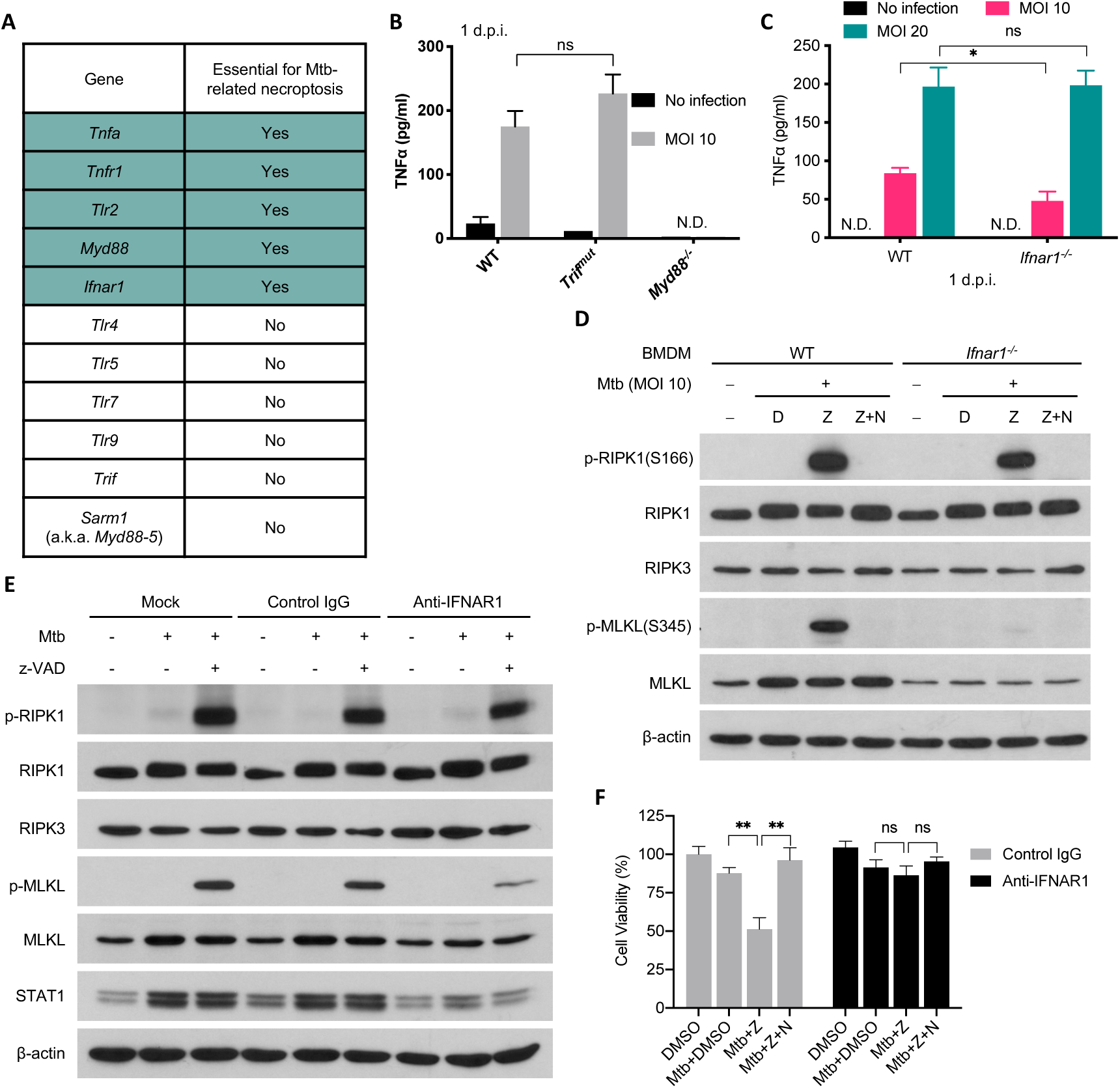
Mtb-related necroptosis relies on TLR-TNF signaling as well as type I IFN signaling. (A) Effects of various gene deficiencies on Mtb-related macrophage necroptosis based on cell viability assay in the presence of Mtb + z-VAD. BMDMs from WT and indicated knockout mice were infected with Mtb in the presence of z-VAD as in Figure 1 at an MOI of 20 and cell viability was assayed 1 d.p.i. by measuring ATP level. Detailed data shown in Figure S2. (B and C) BMDMs were infected with Mtb at an MOI of 10 and cell culture supernatant was harvested 24 h.p.i. and analyzed by TNFα ELISA. (D) Effects of Ifnar1 deficiency on Mtb-related necroptosis activation. BMDMs were treated as in (A) and cells were harvested at indicated time for western blots. (e and f) Effects of blockade of type I IFN signaling on macrophage necroptosis. BMDMs are pretreated with indicated antibody at 20 μg/mL and compounds as in (A) 2 h before infection and the treatment continued till harvest at 10 h.p.i (E) or 1 d.p.i. Shown are immunoblots of cell lysates (E) and cell viability measured by ATP assay (f). Data are means ± SD of three technical replicates (B, C and F) and representative of two experiments. D, DMSO; Z, z-VAD; N, Nec-1; N.D., not detected; ns, not significant; *, p < 0.05; **, p < 0.01 (two-tailed unpaired Student’s *t* test).

Without pathogen infection, in some cell lines, the combination of TNFα and z-VAD induces necroptosis(21, 25), but this was not the case with macrophages in our studies (Figure S3A). Thus, the TNFα produced in response to Mtb is not the only macrophage response to Mtb that is involved in their necroptotic death.

In contrast, type I IFN, another product of Mtb-infected macrophages, was required for their necroptotic death. Absence of *Ifnar1* did not affect TNFα production (Figure 2C) but made the macrophages resistant to Mtb-related, z-VAD-dependent necroptosis (Figure 2A and S2I). *Ifnar1*^−/−^ BMDMs showed a similar level of phosphorylation of RIPK1 as WT cells but less phosphorylated MLKL and less total MLKL, because upregulation of MLKL expression was abrogated in *Ifnar1*^−/−^ BMDMs (Figure 2D), consistent with reports that *Mlkl* is an interferon-stimulated gene (ISG)(15, 26). Blockage of type I IFN signaling starting 2 h before infection with anti-IFNAR1 mAb prevented the upregulation of MLKL and STAT1, another ISG, seen in wild type macrophages, without affecting the basal level of MLKL and STAT1 (Figure 2E). Anti-IFNAR1 mAb did not affect RIPK1 phosphorylation but led to decreased MLKL phosphorylation (Figure 2E) and significant protection from Mtb-related, z-VAD-dependent necroptosis (Figure 2F), suggesting the acute induction of MLKL contributes to the necroptosis in this setting.

Although Mtb infection of macrophages induced the production of TNFα and type I IFNs and both were essential for induction of Mtb-related, z-VAD-dependent necroptosis, the combination of TNFα, IFN-β and/or IFN-α as well as z-VAD did not induce necroptosis of uninfected BMDMs (Figure S3B). Thus, the contribution of Mtb to z-VAD-dependent necroptosis of the macrophages in these studies includes at least one more action or induces one more response beyond induction of TNFα and type I IFNs.

### Mtb-related, z-VAD-dependent necroptosis in a human macrophage cell line

Mtb infection combined with z-VAD treatment resulted in faster cell death of the human U937 macrophages cell line than seen with Mtb infection alone (Figure 3A) and led to phosphorylation of human MLKL (Figure 3B). Nec-1 and GSK’872 abolished cell death (Figure 3A) and MLKL phosphorylation (Figure 3B). The human MLKL inhibitor necrosulfonamide (NSA) inhibited cell death (Figure 3B) but not MLKL phosphorylation (Figure 3A). This is consistent with reports that NSA interacts with the MLKL N-terminal domain without affecting MLKL phosphorylation and this is sufficient to block translocation of phosphorylated MLKL to the membrane(4).

**Figure 3.**
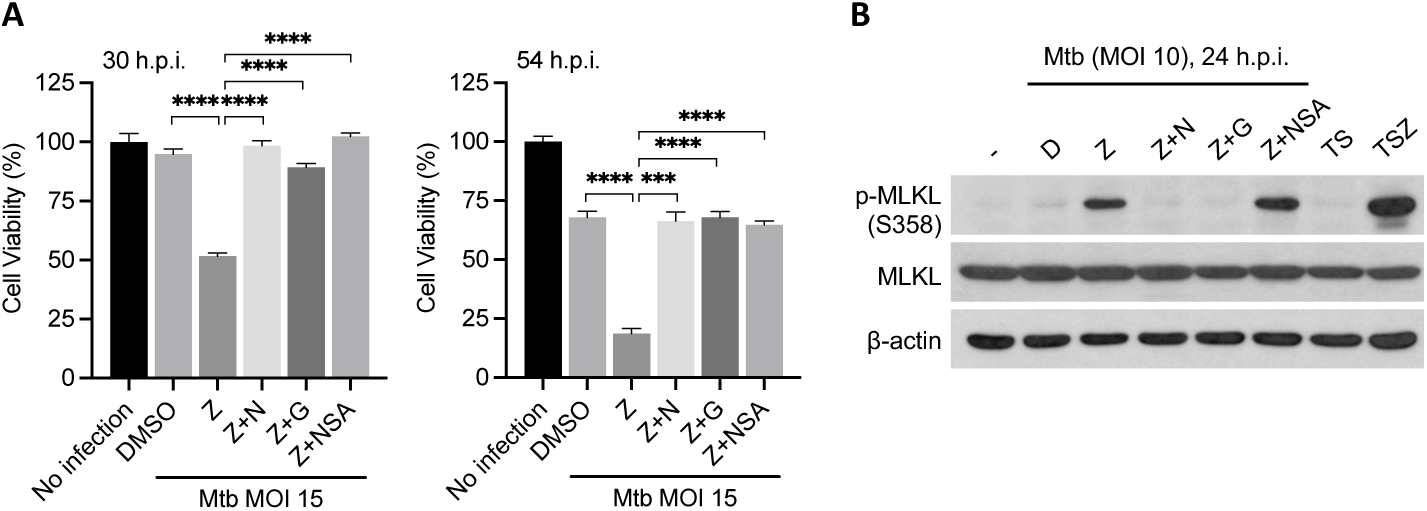
Mtb-related necroptosis is conserved on human macrophages. Human U937 cells were differentiated with 10 ng/mL PMA one day before infection. 2 h before infection, cells were replaced with regular media without PMA and pretreated with compounds at half of the indicated concentration for 2 h before infection of Mtb and then indicated concentration (20 μM except that NSA is used at 3 μM) after washing Mtb away till harvest. (A) Effects of necroptosis inhibitors on Mtb+z-VAD induced U937 cell death. Cell viability measured by ATP assay 30 h.p.i. (left) and 54 h.p.i. (right). (B) Effects of necroptosis inhibitors on Mtb+z-VAD-triggered MLKL phosphorylation on U937 cells. Shown are immunoblots of cell lysates harvest at 24 h.p.i. Data are means ± SD of three technical replicates (A) and representative of two experiments. D, DMSO; Z, z-VAD; N, Nec-1; G, GSK’872; T, TNFα; S, Smac mimetic birinapant; ***, p < 0.001; ****, p< 0.0001 (two-tailed unpaired Student’s *t* test).

### z-VAD-dependent necroptosis is also present in macrophages infected with certain other Gram-positive bacteria

Infection of mouse BMDMs by *Escherichia coli* strain LF82 or *Citrobacter rodentium* in the presence of z-VAD induced a form of cell death that could be blocked by Nec-1, indicating it was likely necroptosis(27). In addition to Mtb, we wondered whether necroptosis also happens to macrophages infected with other mycobacteria or Gram-positive bacteria. Infection of BMDMs with *M. kansasii,* the most pathogenic nontuberculous mycobacterium(28, 29), also resulted in robust necroptosis dependent on *Mlkl* in the presence of z-VAD but not Q-VD-OPH, while *M. abscessus* or *M. marinum* combined with z-VAD were not able to induce necroptosis (Figure 4A). Nec-1 blocked cell death induced by *M. kansasii* (Figure 4A), further confirming it being necroptosis and its dependence on RIPK1.

**Figure 4.**
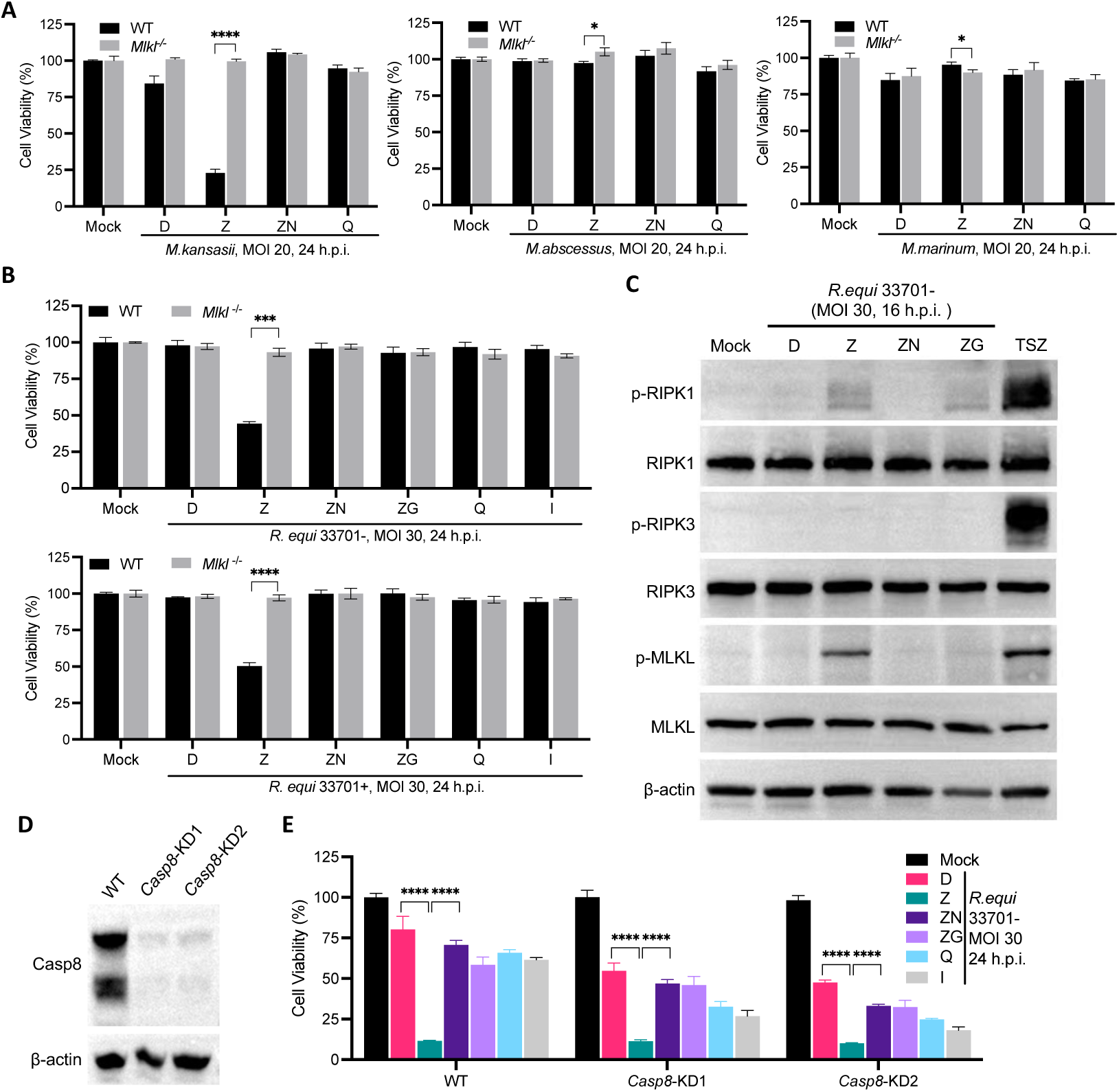
z-VAD-dependent necroptosis is also present in other Gram-positive bacterial infection. (A-C) WT and *Mlkl*^−/−^ BMDMs were pretreated with indicated compounds at half of the indicated concentration for 2 h before pathogen infection and then indicated concentration (20 μM except that GSK’872 is used at 3 μM). (A and B) Cell viability was measured by ATP assay at indicated time. (C) Cells were harvested at 16 h.p.i for western blots. Shown are immunblots of cell lysates. (D and E) Caspase-8 knockdown in J774 did not inhibit *R.equi*-related necroptosis. Validation of Caspase-8 knockdown efficiency by western blots (D) and cell viability after *R.equi* infection at an MOI of 30 measured by ATP assay 24 h.p.i (E). Compound concentrations are the same as in (A-C). Data are means ± SD of three technical replicates (A, B and E) and representative of at least two experiments. D, DMSO; Z, z-VAD; N, Nec-1; G, GSK’872; Q, Q-VD-OPH; I, IETD; *, p < 0.05; ***, p < 0.001; ****, p< 0.0001 (two-tailed unpaired Student’s *t* test).

*R. equi*, a Gram-positive opportunistic pathogen of humans, is evolutionarily related to Mtb and primarily infects and replicates in alveolar macrophages(30). It was recently reported that *R. equi* also induces a type I IFN response dependent on host cGAS-STING pathway(31). Indeed, *R. equi* infection of BMDMs elicited upregulated expression of type I IFNs, downstream ISGs and other inflammatory cytokines including TNFα, independent of its virulence plasmid (Figure S4A). Therefore, we speculated whether *R. equi*-infected macrophages are predisposed to necroptosis. Like Mtb, *R. equi*-infected BMDMs underwent fast cell death in the presence of z-VAD but not Q-VD-OPH or IETD and cell death could be blocked by Nec-1 or GSK’872 (Figure 4B). Furthermore, deficiency of MLKL in BMDMs afforded complete protection (Figure 4B). In addition, z-VAD-hastened cell death was seen in macrophages infected with either avirulent (33701−) or virulent (33701+) *R. equi* (Figure 4B), consistent with the expression of type I IFNs, ISGs and TNFα not relying on its virulent plasmid.

The mouse macrophage cell line J774A.1 underwent *R. equi*-related, z-VAD-dependent necroptosis similarly to BMDMs (Figure S4B). *R. equi* + z-VAD robustly induced phosphorylation of RIPK1 and MLKL (Ser345), but not phosphorylation of RIPK3 at Thr231 and Ser232 in J774A.1 cells (Figure 4C), echoing what we saw in BMDMs infected with Mtb and treated with z-VAD.

As caspase-8 is essential for cell survival, we were not able to completely knock out the *Casp8* gene in J774A.1 cells. Instead, we used macrophages haplo-deficient in *Casp8*, which had drastically reduced expression of caspase-8 protein (Figure 4D). The haplo-deficient cells showed pronounced sensitivity to *R. equi* infection; however, the addition of z-VAD still resulted in more cell death than in the DMSO control, which could be reverted to a similar level by Nec-1 and GSK’872 (Figure 4E). These data further confirm that z-VAD is very likely to act on non-caspase targets in macrophages undergoing both Mtb- and *R. equi*-related necroptosis.

### Necroptosis appears to contribute to disease pathology in *Sp140*^−/−^ mice infected with a high Mtb burden

We then confirmed and extended the results of Stutz et al.(15) in our studies of *Mlkl*^+/+^ and *Mlkl*^−/−^ mice on the C57BL/6 background infected with 200-300 CFU of Mtb by inhalation. Starting from day 28 post infection, we observed an increase in the MLKL protein level in *Mlkl*^+/+^ lung homogenates and not in *Mlkl*^−/−^ mice, but we did not detect phosphorylation of MLKL (Figure 5A). Moreover, there was no significant difference between *Mlkl*^+/+^ and *Mlkl*^−/−^ mice with respect to the Mtb burden in lungs, spleens and livers (Figure 5B).

**Figure 5.**
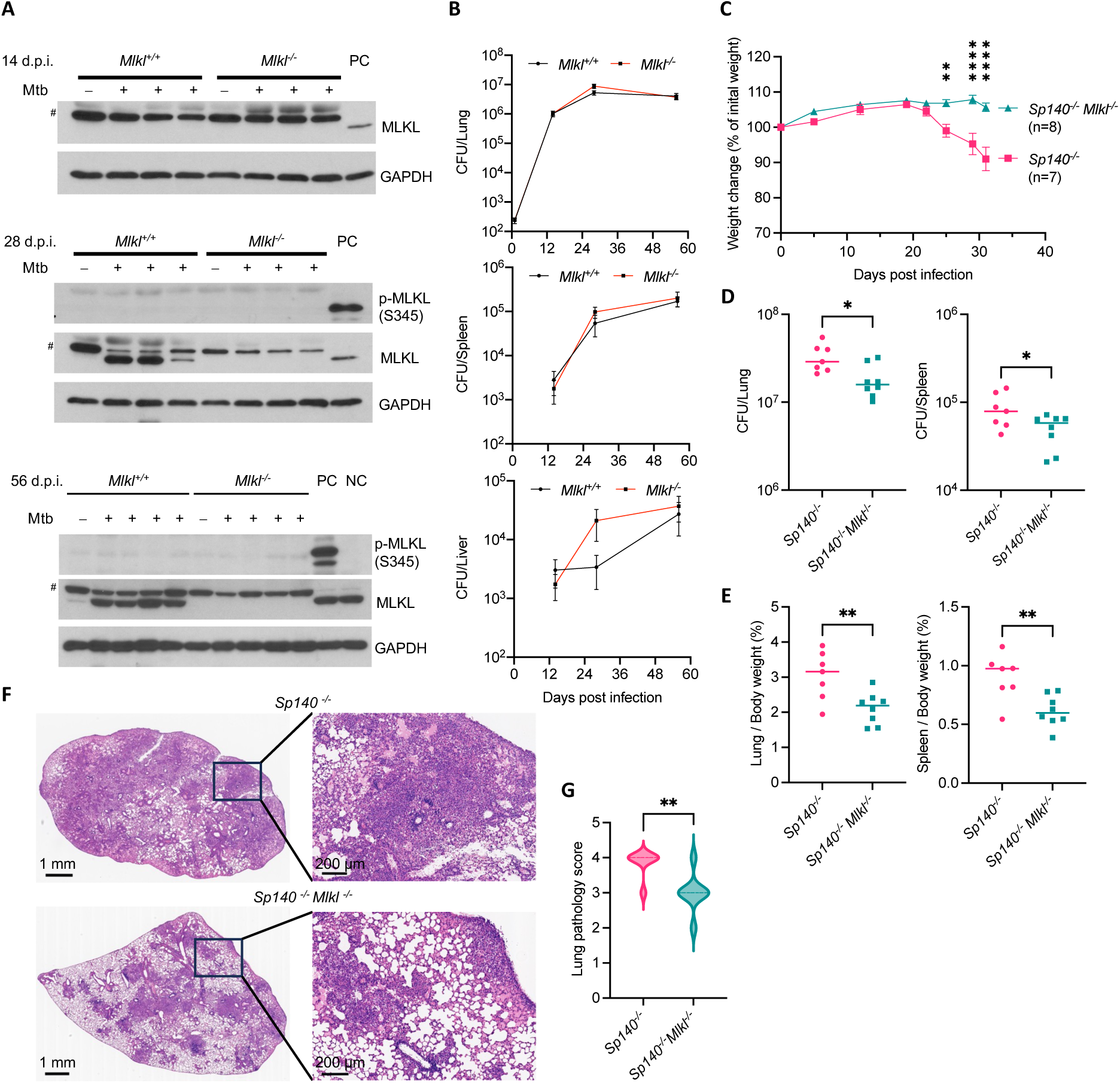
The absence of MLKL delays TB progression in Mtb-infected *Sp140*^−/−^ mice. (A and B) *Mlkl*^+/+^ and *Mlkl*^−/−^ mice were infected with 200-300 CFU of Mtb on day 0. (A) Mtb infection on mice resulted in MLKL upregulation but not MLKL phosphorylation in lung homogenates. Samples were prepared from uninfected and infected mice harvested on indicated days post infection (d.p.i.). Each lane represents one mouse. Shown are immunoblots. PC, positive control, Mtb + z-VAD -treated BMDM lysates; NC, negative control, mock-treated BMDM lysates. #, non-specific band. (B) Mtb burdens in multiple organs of *Mlkl*^+/+^ and *Mlkl*^−/−^ mice. n=2 for each strain on day 0, n=5∼8 for each strain on day 14, 28 and 56. (C-G) *Sp140*^−/−^ and *Sp140*^−/−^Mlkl^−/−^ mice were infected with 4000-6000 CFU of Mtb on day 0. For (D-G), mice were euthanized on day 31. (C) Weight changes of Mtb-infected mice. (D) Mtb burdens in lungs (left) and spleens (right) of mice. (E) The ratio of lung (left) or spleen (right) weight to whole body weight in Mtb-infected mice. (F) Representative HE staining of lung sections of Mtb-infected mice. (G) Pathology scores of lung sections of Mtb-infected mice. n=7 for *Sp140*^−/^ mice and n=8 for *Sp140*^−/−^*Mlkl*^−/−^ mice. Data are means ± SEM (B-E and G). *, p < 0.05; **, p < 0.01; ****, p< 0.0001 (two-tailed unpaired Student’s *t* test).

Given that *Mlkl* is an ISG and previous reports showed that *Sp140*^−/−^ mice had elevated IFN-I levels after Mtb infection due to abrogation of negative regulation of *Ifnb1* mRNA by SP140 (32, 33), we wondered whether the deficiency of SP140 would potentiate the induction of *Mlkl* and affect TB disease progression in mice. We infected *Sp140*^−/−^ and *Sp140*^−/−^ *Mlkl*^−/−^ mice on the C57BL/6 background with 400-500 CFU of Mtb by inhalation. Due to the relatively low virulence of this particular strain, the bacterial burden in lungs of *Sp140*^−/−^ mice reached only around 10^5^ CFU and additional deficiency of *Mlkl* did not result in significant weight change nor affect Mtb burdens in lungs, spleens or livers (Figure S5A-B). The ratio of lung weight/whole body weight was lower in Mtb-infected *Sp140*^−/−^ *Mlkl*^−/−^ mice than in Mtb-infected *Sp140*^−/−^ mice, yet the lung pathology and spleen/body weight ratios were similar in both *Sp140*^−/−^ and *Sp140*^−/−^ *Mlkl*^−/−^ mice (Figure S5C-E).

However, when mice were infected with an Mtb inoculum high enough to cause weight decrease in *Sp140*^−/−^ mice, *Mlkl* deficiency in the *Sp140*^−/−^ background blocked the decrease of weight significantly and also reduced Mtb burdens in lungs and spleen (Figure 5C-D), leading to less inflammation in both organs as judged by organ/whole body weight ratio as well as by hematoxylin and eosin staining of infected lungs (Figures 5E-G). The impact of *Mlkl* deficiency suggests that necroptosis contributes to TB pathology in TB-susceptible *Sp140*^−/−^ mice.

## Discussion

Mtb infection, in concert with the type I IFN that the infection induces, leads to death of mouse macrophages, at least in part through the impact of host ACOD1 on lysosomal membrane permeabilization(19, 34). The present work establishes that there is a point of control in macrophages whose engagement switches the cell death mechanism to necroptosis in a manner that is also dependent on type I IFN, additionally dependent on TNFα induced by the infection, and further dependent on at least one more contribution of Mtb or the macrophage response to Mtb. The existence of this control point was revealed by a non-canonical effect of z-VAD.

What we observed is not restricted to infection of macrophages by Mtb. Infection of mouse BMDMs by *Escherichia coli* strain LF82 or *Citrobacter rodentium* in the presence of z-VAD induced a form of cell death that could be blocked by Nec-1, indicating it was likely necroptosis(27). Cell death in this setting relied on both TNF signaling and type I IFN signaling, similar to the necroptosis triggered by Mtb infection in macrophages exposed to z-VAD. We further identified that other Gram-positive pathogens including *M. kansasii* and *R. equi* also triggered macrophage necroptosis in the presence of z-VAD.

The canonical function of z-VAD is to inhibit caspases in a cell; its ability to promote necroptosis has been ascribed to its inhibition of caspase-8. Many bacteria and viruses are equipped with machinery to prevent caspase activation(35–41). Concurrent infection of macrophages with Mtb and other bacteria or viruses could theoretically lead to necroptosis through caspase inhibition. However, in our studies, caspase inhibition was insufficient to switch pathogen-infected macrophages to a necroptotic death pathway, based on the following observations: (1) We previously found that Mtb infection alone strongly inhibits the activity of caspase-8 and caspase-3 and does not lead to necroptosis(19). (2) In contrast to z-VAD, the more specific caspase-8 inhibitor IETD and the more specific and potent pan-caspase inhibitor Q-VD-OPH did not sensitize pathogen-infected macrophages to necroptosis. (3) TSZ triggered much faster necroptosis in macrophages than TSQ. (4) z-VAD is known to inhibit other proteases besides caspases(42–45). (5) z-VAD still triggers fast necroptosis in casp8-KD J774A.1 macrophages upon *R. equi* infection. We speculate that z-VAD may target protein(s) that are expressed in macrophages but not in the cell lines often used for studies of necroptosis. Cell type-specific necroptotic signaling has been observed in the differential sensitivity to TNFα on the part of FADD-deficient MEF cells and Jurkat cells(6).

Our studies also highlighted the potential of RIPK3 to undergo different patterns of phosphorylation or other post-translational modification to sustain necroptosis. Many of the serine and threonine residues of RIPK3 are subject to phosphorylation(6, 46). Among them, phosphorylation of human RIPK3 on Ser227 and murine RIPK3 (mRIPK3) on Thr231 and Ser232 is thought to be essential for binding MLKL in a species-specific way to mediate necroptosis induced by TNF signaling and by LPS + z-VAD(3, 46, 47). Surprisingly, we found that Mtb/*R.equi*-related, z-VAD-dependent necroptosis did not lead to detectable phosphorylation of mRIPK3 on Thr231 and Ser232. Nonetheless, necroptosis depended on the presence of RIPK3, was blocked by the RIPK3 inhibitor GSK’872 and was accompanied by phosphorylation of the RIPK3 substrate mMLKL on Ser345, which in turn was blocked by the RIPK3 inhibitor GSK’872. These results strongly suggest that RIPK3 was activated, but most likely accompanied with phosphorylation on residues other than Thr231 and Ser232 or with other types of post-translational modification to recruit and phosphorylate MLKL. This deviation from the classical paradigm for necroptosis suggests that the pathway is more plastic than had been appreciated.

Studies on necroptosis induced by LPS plus z-VAD or classical TSZ stimuli suggested that subtle changes in MLKL protein levels can dramatically affect cell sensitivity to necroptotic stimuli and that constitutive interferon signaling maintains cytosolic MLKL above a certain threshold to license necroptosis(48). In addition to this, we found that the role of type I IFN signaling in the Mtb-related, z-VAD-dependent necroptosis of macrophages includes an acute increase in expression of MLKL upon Mtb infection.

Results of our experiments with Mtb infection of *Mlkl*^−/−^ mice are in agreement with previous studies that MLKL-dependent necroptosis is unlikely to account for pathology in Mtb-infected WT B6 mice(15, 16). However, WT B6 mice rarely develop necrotic pulmonary lesions after Mtb infection(49), which might obscure the possibility of identifying the presence of necroptosis in affected lungs. Deficiency of SP140 in B6 mice results in higher type I IFN level and phenocopies the response of 129S2 and C3HeB/FeJ strains to Mtb infection with respect to formation of necrotic granulomas as seen in some forms of TB in humans (49, 50). Indeed, compared to *Sp140*^−/−^ mice, *Sp140*^−/−^ *Mlkl*^−/−^ mice showed less weight loss, lower Mtb burdens and alleviated lung pathology after high-dose Mtb infection, possibly due to less release of inflammatory material from dying cells. Taken together, our findings suggest that necroptosis can contribute to TB disease under some circumstances. Identification of the relevant targets of z-VAD may provide more insight into how necroptosis is regulated during TB.

## Acknowledgements

We thank E. Pamer (then at Memorial Sloan Kettering Cancer Center) for *Myd88*^−/−^ and *Tlr4*^−/−^ mice, A. Choi (Weill Cornell Medicine) for providing *Mlkl*^−/−^ mice, and R. Vance (University of California, Berkeley) for providing *Sp140*^−/−^ mice. We thank T. Dong (Southern University of Science and Technology, China) for *M. marinum* strain, R. Waston and N. Cohen (Texas A&M University) for *R. equi* 33701+ and 33701-strains. We thank the Biosafety Level 3 Laboratory of Shenzhen University for their support during this research. For help with studies that were not conclusive enough to include, we thank V. Dartois at Hackensack Meridian Center for Discovery and Innovation for sections from tuberculous human lungs, J. Wolchock and T. Merghoub at Memorial Sloan Kettering Cancer Center for sections from human melanoma; N. Altorki (Weill Cornell Medicine) for sections of normal lung from subjects with lung cancer; B. Chen (Weill Cornell Medicine) for lung sections from autopsy samples; S. Monette, A. Piersigilli, M. S. Jiao (Memorial Sloan Kettering Cancer Center) and the Core Facility, University of Pittsburgh Medical Center) for immunohistochemistry; and Y. Chen (Weill Cornell Medicine) for advice in the interpretation of immunohistochemistry. We thank members of the Nathan laboratory and Zhang laboratory for discussions and technical assistance. This work was supported by the National Natural Science Foundation of China (Grant No.32470183 to L.Z.), Guangdong Basic and Applied Basic Research Foundation (Grant No.2025A1515010652 to L.Z.), the National Institutes of Health (Grant No. 1R01AI138940 to C. N, P30CA047904 and DP2GM146320 to Y.-N.G.) and the Milstein Program in Chemical Biology and Translational Medicine (to C. N). The Department of Microbiology and Immunology of Weill Cornell Medicine is supported by the William Randolph Hearst Foundation.

## Author contributions

L. Z. made the initial observation and conceived the study; R. H. performed experiments assisted by P. L. and J. Han. X. J. contributed to mouse infection experiments; J. Huang, G. H. and D.P. provided technical assistance; G. L. supervised the wok of J. Huang and G. H.; Y.-N.G. contributed to immunohistochemistry analysis; L. Z. and C. N. supervised the project; R.H., L. Z. and C. N. analyzed the data and wrote the manuscript with input from other authors.

## Materials and methods

### Cell lines and primary macrophages

Human U937 cells (ATCC, Cat# CRL-1593.2) were cultured in RPMI 1640 supplemented with 10% fetal bovine serum (FBS), 2 mM L-glutamine, 10 mM HEPES pH 7.5 and 1 mM sodium pyruvate and differentiated with 10 ng/mL phorbol 12-myristate 13-acetate (PMA) one day before infection. L929 (ATCC, Cat# CCL-1) and J774A.1 (ATCC, Cat# TIB-67) were cultured in Dulbecco’s modified Eagle’s medium (DMEM) supplemented with 10% FBS, 2 mM L-glutamine, 10 mM HEPES pH 7.5 and 1 mM sodium pyruvate. L929-conditioned medium (LCM) and bone marrow-derived macrophages (BMDMs) were prepared as previously described(19). All cells were cultured at 37 °C in a 5% CO_2_ incubator unless otherwise specified.

### Mice

WT C57BL/6 (#000664), WT C57BL/6N (#005304), *Ripk3*^mut^ (#025738)*, Tnfα*^−/−^ (#005540), *Tnfr1*^−/−^ (#003242), *Tlr2*^−/−^(#004650), *Trif*^mut^(#005037), *Tlr5*^−/−^(#008377), *Tlr7*^−/−^ (#008380), *Tlr9*^−/−^(#34329-JAX), *Sarm1*^−/−^(#018069) and Cas9 knockin mice (#026179) (all on the C57BL/6 background) were purchased from Jackson Laboratory. Cas9-het mice were prepared and used as previously described(19). *Mlkl*^+/+^ and *Mlkl*^−/−^ mice were bred in-house from a heterozygote X heterozygote mating scheme and genotyped before infection. Mice of both sexes aged 6∼12 weeks were used for Mtb infection experiments. At most time points, the mice harvested from different experimental groups were sex- and age-matched. All mice were housed in a specific pathogen-free facility or a BSL3 vivarium if infected by Mtb. All mouse experiments were approved by and performed in accordance with requirements of the Institutional Animal Care and Use Committees at Weill Cornell Medicine or Southern University of Science and Technology.

### Reagents

Mouse TNFα (#T0157), Nec-1 (#N9037) and were purchased from Sigma. Mouse IFN-αA (#12100-1) and mouse IFN-β (#12405-1) were from PBL Assay Science. z-VAD-FMK (#A1902) were from Apexbio. Smac mimetic birinapant (#501015095), GSK’872 (#5303890001) and z-IETD-FMK (#FMK007) were from Fisher Scientific. Necrosulfonamide (#20844), Q-VD-OPH (#15260) and Ac-YVAD-CMK (#10014) were from Cayman Chemical. Mouse monoclonal anti-human IFN-γ Rα chain (as control IgG; #G737) and mouse monoclonal anti-mouse IFNAR1 (MAR1-5A3; #I-1188) were purchased from Leinco Technologies. Antibodies for β-actin (#sc-47778) and RIPK3 (#sc-374639) were from Santa Cruz Biotechnology. Antibodies for phospho-RIP1 (Ser166; #31122S), MLKL (#37705S for mouse, #14993S for human), phospho-MLKL (Ser345; #37333S) and STAT1 (#9172S) were from Cell Signaling Technology. Phospho-RIPK3 (Thr231/Ser232; #ab222320) antibody was from Abcam and RIPK1 antibody (#610459) from BD Biosciences. Anti-GAPDH (#G041) was purchased from Abm.

### Bacterial culture and macrophage infection

Mycobacteria were grown in Middlebrook 7H9 medium supplemented with 0.5% glycerol, 10% oleic acid–dextrose–catalase (BD Biosciences; #212351) and 0.02% tyloxapol (7H9 complete medium). *Mycobacterium tuberculosis* (Mtb), *M. kansasii* and *M. marinum* were cultured in a 5% CO_2_ incubator at either 37 °C (Mtb and *M. kansasii)* or 32 °C (*M. marinum*). *M. abscessus* was grown with constant shaking at 37 °C. Mycobacterial single cells were prepared as previously described(19) and added to macrophages to initiate infection. After infection for 4 h (Mtb), 3 h (*M.kansasii*) or 1 h (*M. marinum* and *M. abscessus*), macrophages were washed twice with warm PBS and then fresh medium was replaced. No antibiotics were used in the preparation, infection or post-infection incubation of macrophages.

For *Rhodococcus equi* (*R.equi*) infection, one colony of *R.equi* was inoculated into 5 ml of brain-heart infusion broth (Thermo Scientific Oxoid; CM1135) and shaken overnight 30 °C, then indicated culture was subcultured into fresh BHI and shaken at 37 °C for 3-4 h until the optical density of 600 nm (OD 600) reached to 0.3-0.5. The bacteria were washed twice with PBS and resuspended in DMEM complete medium, then added to macrophages. 1 h later, macrophages were washed twice with warm PBS and medium containing 100 μg/ml gentamycin was replaced to kill extracellular *R.equi*. 1 h later, fresh medium containing 10 μg/ml gentamycin was replaced after macrophages were washed with warm PBS.

For western blot, macrophages were harvested at the indicated time point and lysed with 1 X SDS loading buffer at 95 °C for 30 min. For quantitative PCR, total RNA was extracted using EZ-press RNA Purification Kit (EZBioscience; B0004D). cDNA was synthesized with Hifair® III 1st Strand cDNA Synthesis Kit (Yeasen; 11141ES10) according to the manufacturers’ instructions. qRT-PCR was performed using ChamQ Universal SYBR qPCR Master Mix (Vazyme; Q711). Data were analyzed on CFX96 Touch Real-Time PCR Detection System (Bio-Rad).

### Cell viability assays

Viability of BMDMs was assayed by measuring ATP levels using a Promega CellTiter-Glo kit according to the manufacturer’s instructions. Percentage viability was normalized to that of uninfected cells.

### ELISA

Cell culture supernatant was centrifuged at 21,130 g at 4 °C for 5 min and then passed through a 0.22 μm cellulose acetate filter before being removed from the BSL3 facility. Samples were diluted 3-to 4-fold before ELISA were performed in accordance with the manufacturers’ instructions.

### Aerosol infection of mice and organ harvest

Mtb infection of mice was carried out as previously described(19). Briefly, log phase cultures of Mtb H37Rv strain grown in Middlebrook 7H9 medium containing 0.5% glycerol, 10% albumin–dextrose–saline (ADN) supplement and 0.05% Tween-80 were used to prepare a suspension enriched in single cells. The required volume was centrifuged for 10 min at 3,082 g and the pellet resuspended in PBS. Mice were infected using a Glas-Col Inhalation Exposure System using the Mtb inoculum prepared as above. On day 1 post infection, 2 mice were euthanized and lung homogenates in PBS were plated on 7H10 agar supplemented with 0.5% glycerol and 10% OADC to determine the initial bacterial load. On indicated days, mice were euthanized with CO_2_ and lungs, livers and spleens were harvested. Each organ (except the upper lobe of the left lung) was homogenized in PBS (if the homogenates were to be used for western blot, the PBS was supplemented with 1% Roche EDTA-free protease inhibitor cocktail and PhosStop), serially diluted and plated on 7H10 agar for determination of CFU at 3 weeks. The upper lobes of the left lungs were fixed in 10% formalin.

For western blot, lung homogenates were centrifuged at 21,130 g at 4 °C for 20 min. Supernatant was centrifuged again and passed through a 0.22 μm cellulose acetate filter before being removed from the BSL3 facility. The protein concentration was quantified by a BIO-RAD *DC*™ Protein Assay kit and ∼50-60 μg protein used for SDS-PAGE and western blot.

For histopathological analysis, formalin-fixed lungs were embedded in paraffin, sectioned and then stained with Hematoxylin and Eosin (H&E). The stained sections were scanned by a digital pathology scanner (KFBIO; KF-PRO-120) and pathological severity was scored in a blinded fashion based on the extent of inflammation: 0, 1, 2, 3, and 4 correspond to 0-4%, 5-20%, 21-40%, 41-60%, and 61-100% of the lung area affected by inflammation, respectively.

### Generation of J774A.1 knock-out cells

All J774A.1 knock-out cell lines were generated as previously described(51). 5 X 10^5^ J774A.1 cells were spin infected (800 rpm, 1.5 h, 37 ℃) with gRNA lentivirus packaged from LentiCRIPSRv2 vectors. 24 h later, fresh medium was replaced and cells were cultured for 24 h before adding 3 μg/ml puromycin for 3 days. Then genomic DNA from live cells was prepared, and gRNA efficiency was validated by SURVEYOR assay. Single cells were then sorted using BD FACS Aria SORP, and the genotype of each cell clone was verified by sequencing of the PCR fragments. The efficiency of specific gene knock-out was validated by western blot. The gRNA sequences are as follow: *Casp8*: 5’-TGAGATCCCCAAATGTAAGC-3’; *Ripk3*-gRNA1: 5’-GTGGGACTTCGTGTCCGGGC-3’; *Ripk3*-gRNA2: 5’-AACCCGAGTGCCCTCGGCCC-3’; *Mlkl*-gRNA1: 5’-GCGTCTAGGAAACCGTGTGC-3’; *Mlkl*-gRNA2: 5’-TGGGCCGTTTTGATGAAGTC-3’.

### Electroporation of guide RNAs into Cas9 BMDMs

LentiGuide plasmids (Addgene, #52963) with the correct gRNA insertion were used as a template to amplify hU6 promoter-gRNA fragments and then electroporate into Cas9 BMDMs as previously described(19). The gRNA sequences used are: negative control (NC); 5’-GAACTCGTTAGGCCGTGAAG-3’ (51); *Mlkl*: 5’-GCACACGGTTTCCTAGACGC-3’.

### Statistical analyses

Results of cell viability assays and ELISAs in macrophages are represented as means ± SD and analyzed by two-tailed unpaired Student’s t test using GraphPad Prism. Mtb burden in mouse organs was represented as means ± SEM. ns, not significant; *, p < 0.05; **, p < 0.01; ***, p < 0.001, ****, p< 0.0001.

---

**Figure S1.**
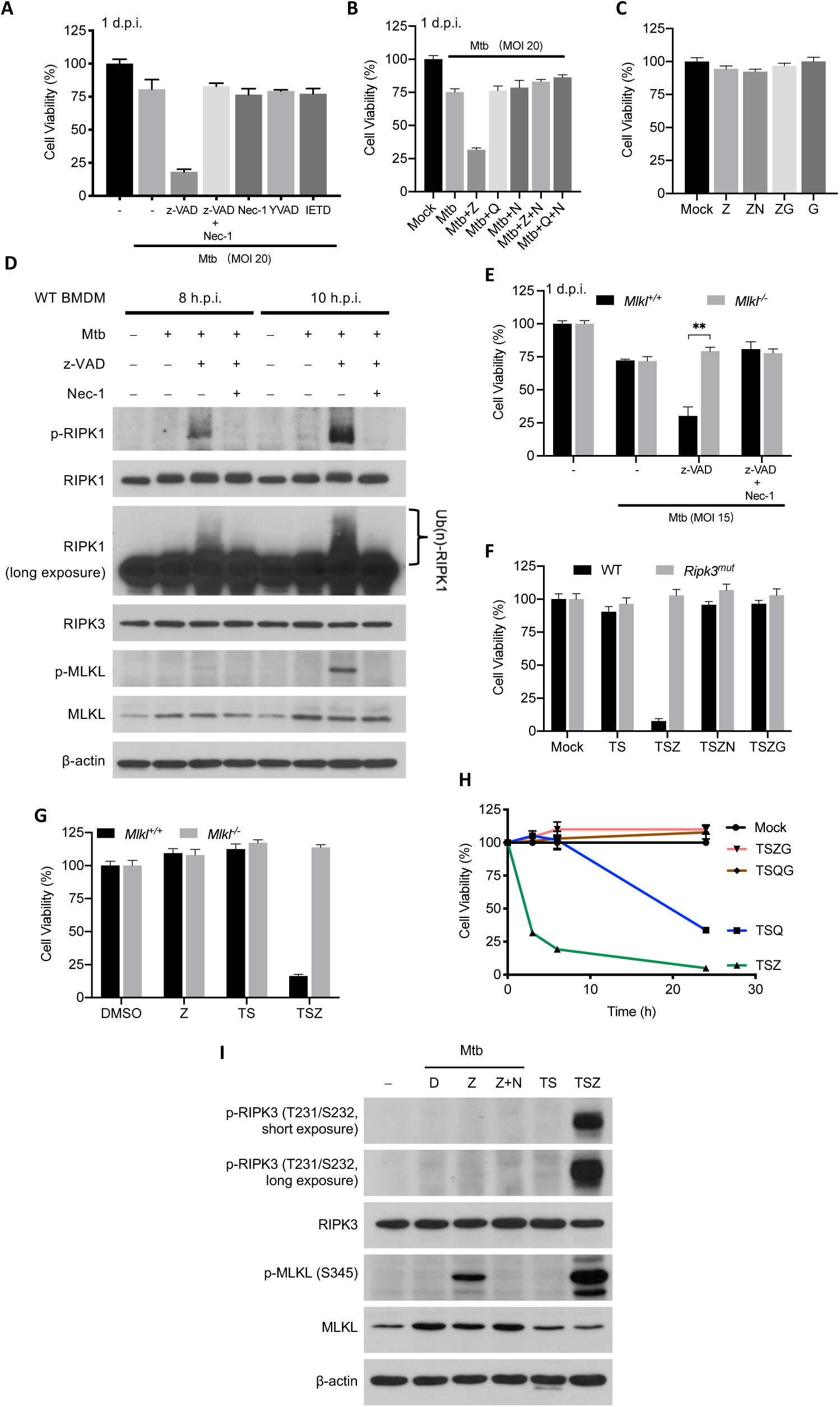
z-VAD sensitizes Mtb-infected BMDMs to necroptosis, which relies on host RIPK3 and MLKL, related to Figure 1. For infection experiment, BMDMs were pretreated with compounds at half of the indicated concentration for 2 h before infection of Mtb and then indicated concentration (20 μM except that GSK’872 is used at 3 μM) after washing Mtb away till harvest. (A and B) Effects of various caspase inhibitors on Mtb-infected macrophages. WT BMDMs were pretreated with indicated compounds and cell viability was measured on 1 d.p.i. (C) z-VAD, Nec-1 and GSK’872 are not cytotoxic without Mtb infection. WT BMDMs were treated with indicated compounds for 24 h and then assayed for cell viability. (D) Mtb + z-VAD induced RIPK1 and MLKL phosphorylation in a time-dependent manner. WT BMDMs were pretreated with indicated compounds 2 h before being infected with Mtb at an MOI of 10. Cell lysates were harvested at indicated time and subjected to immunoblots. (E) MLKL is essential for Mtb + z-VAD-triggered necroptosis. BMDMs were pretreated with indicated compounds 2 h before being infected with Mtb at an MOI of 15. Cells were assayed 1 d.p.i. (f and g) Mutation of *Ripk3* or loss of *Mlkl* in BMDMs renders cells resistant to TSZ-triggered necroptosis. BMDMs were treated with indicated compounds for 4 h and then assayed for cell viability by measuring ATP level. (h) The viability curve of WT BMDMs in response to different necroptotic stimulus. Cell viability was measured by ATP level on 3, 6 and 24 h after treatment. (i) Mtb + z-VAD did not induce classical RIPK3 phosphorylation. Cells were treated, infected and assayed as in (d). TS and TSZ treatment lasted for 2 h. T, TNFα, 10 ng/mL; S, Smac-mimetic birinapant, 10 μM; Z, z-VAD, 20 μM; Q, Q-VD-OPH, 20 μM; G, GSK’872, 3 μM; N, Nec-1, 20 μM. Data shown as means ± SD of three technical replicates (a-c and e-h) and representative of two independent experiments.

**Figure S2.**
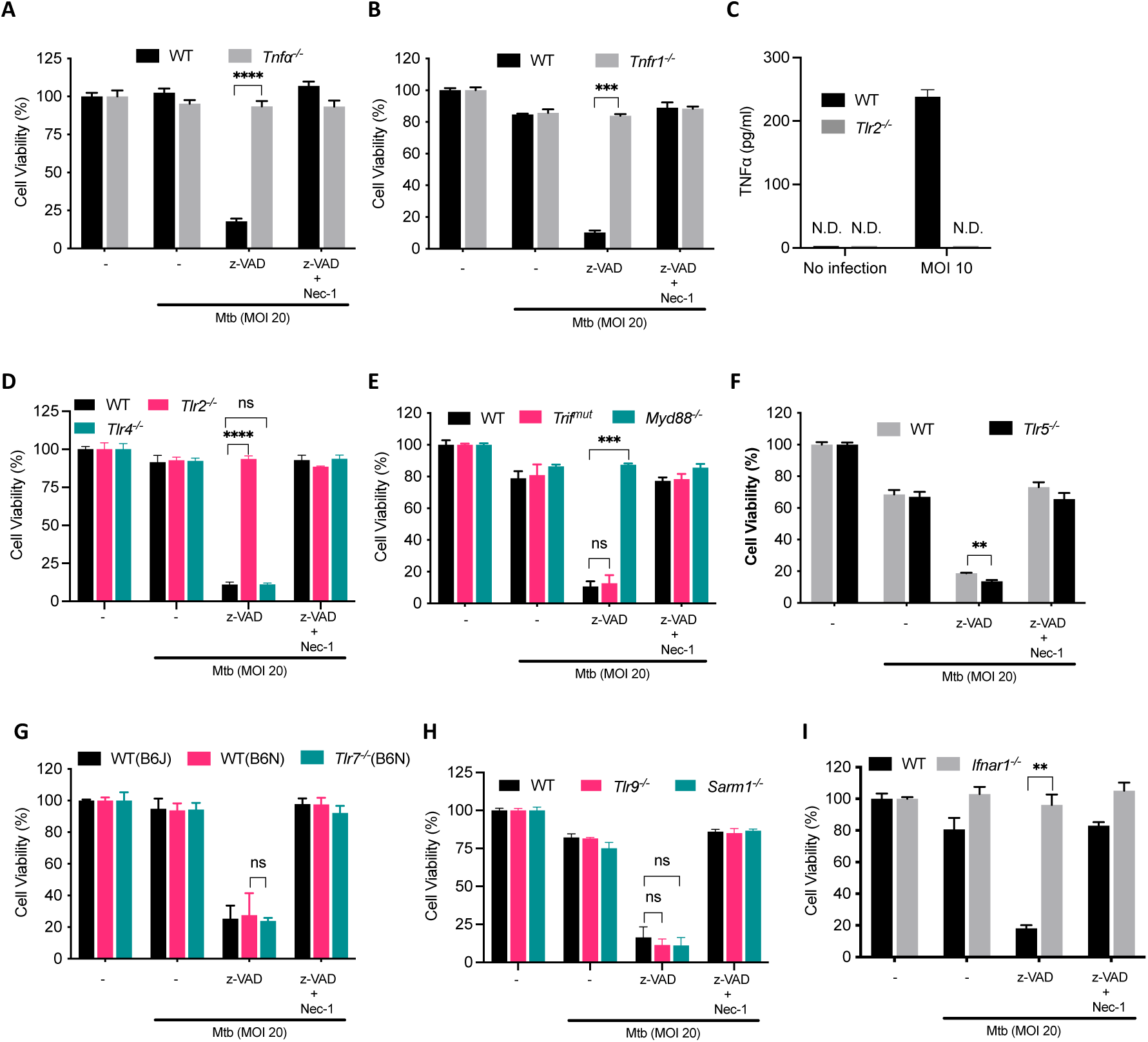
Mtb-related necroptosis depends on TLR2-TNF signaling as well as type I IFN signaling, related to Figure 2. (A-B, D-I) Effects of various gene deficiencies on Mtb-related macrophage necroptosis based on cell viability assay in the presence of Mtb + z-VAD. BMDMs from WT and indicated knockout mice were pretreated compounds with at 10 μM for 2 h before infection of Mtb at an MOI of 20 and then 20 μM after washing Mtb away till harvest. Cell viability was assayed 1 d.p.i. by measuring ATP level. Data are means ± SD of three technical replicates. Data are summarized in Figure 2A. (C) WT and *Tlr2*^−/−^ BMDMs were infected with Mtb and culture supernatant was harvested 24 h.p.i. and measured by TNFα ELISA. N.D., not detected.

**Figure S3.**
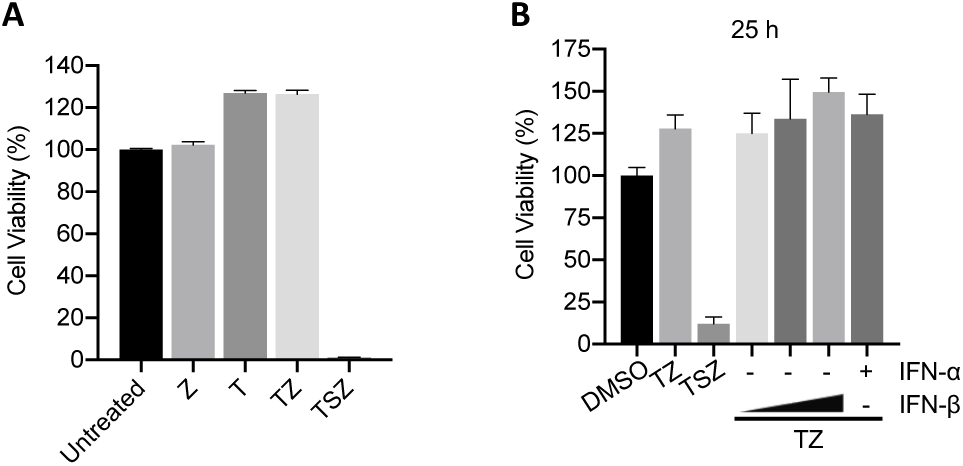
Mtb-induced production of TNFα and type I IFNs are important for necroptosis triggered by Mtb + z-VAD, related to Figure 2. (A) TNFα plus z-VAD does not elicit necroptosis. WT BMDMs were treated with indicated compounds for 31 h and cell viability was analyzed by measuring ATP level. (B) The combination TNFα and type I IFN does not replace Mtb infection in inducing necroptosis. Cells are treated with indicated compounds and cytokine for 25 h and then assayed as in (A). T, TNFα, 10 ng/mL; S, Smac-mimetic birinapant, 10 μM; Z, z-VAD, 20 μM; IFN-α, 1,000 U/ml; IFN-β: 100, 500, and 1,000 U/ml.

**Figure S4.**
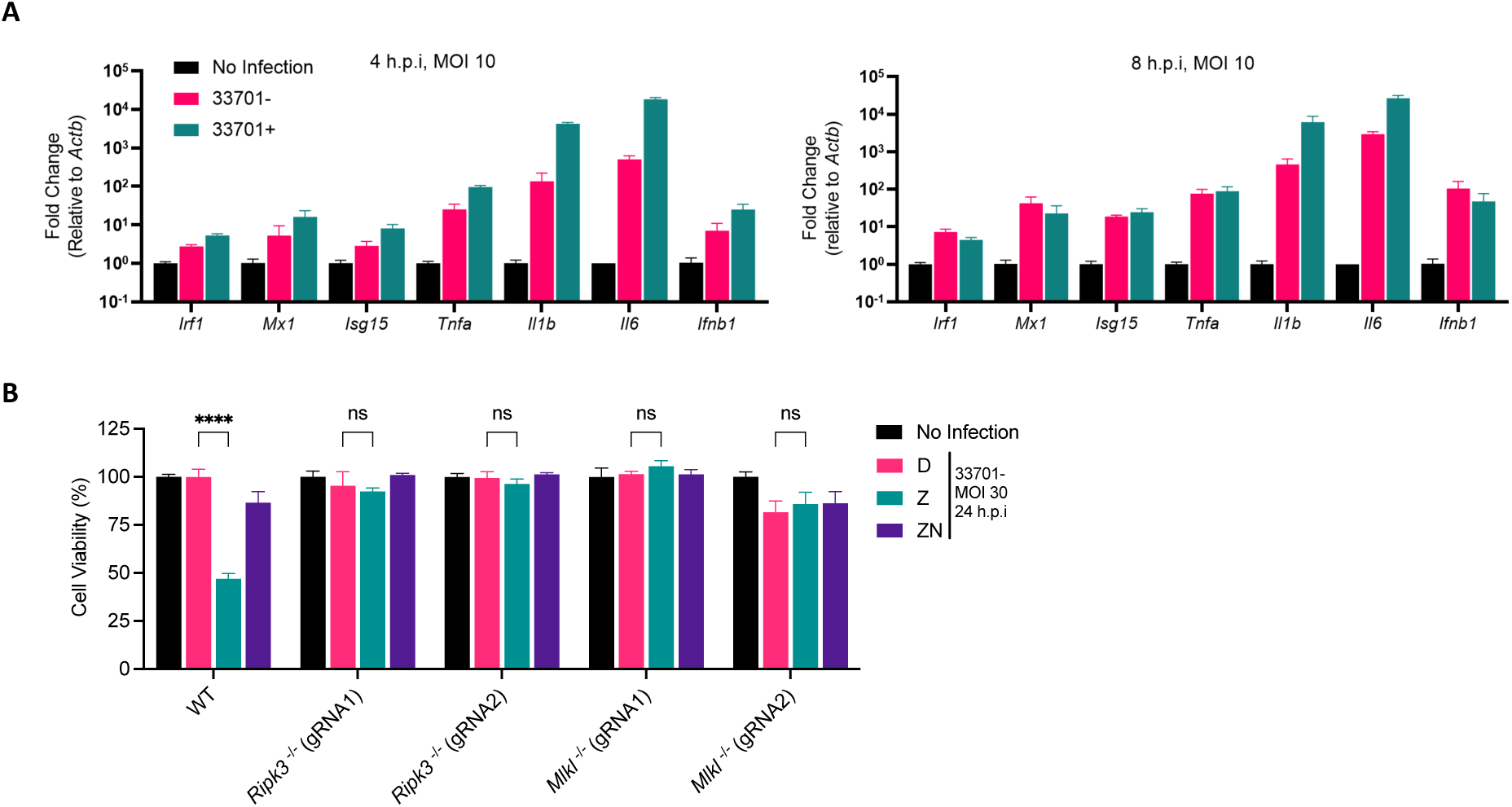
*R.equi* activates type I IFN signaling and induces RIPK3 and MLKL-dependent cell death in the presence of z-VAD, related to Figure 4. (A) qPCR analysis on *R.equi*-infected BMDMs at 4 h.p.i (left) and 8 h.p.i (right). Total RNA was extracted and used for cDNA reverse transcription before qPCR analysis. (B) J774.1 cells were pretreated at half of the indicated concentration for 2 h before *R. equi* 33701-strain infection and then indicated concentration till measuring ATP level 24 h.p.i. KO cells are generated by CRISPR-Cas9–mediated targeting independently using gRNA1 or 2 for both *Ripk3* and *Mlkl*. D, DMSO; Z, z-VAD, 20 μM; N, Nec-1, 20 μM. Data shown as means ± SD of three technical replicates and representative of two independent experiments. ns, not significant; ****, p< 0.0001 (two-tailed unpaired Student’s *t* test).

**Figure S5.**
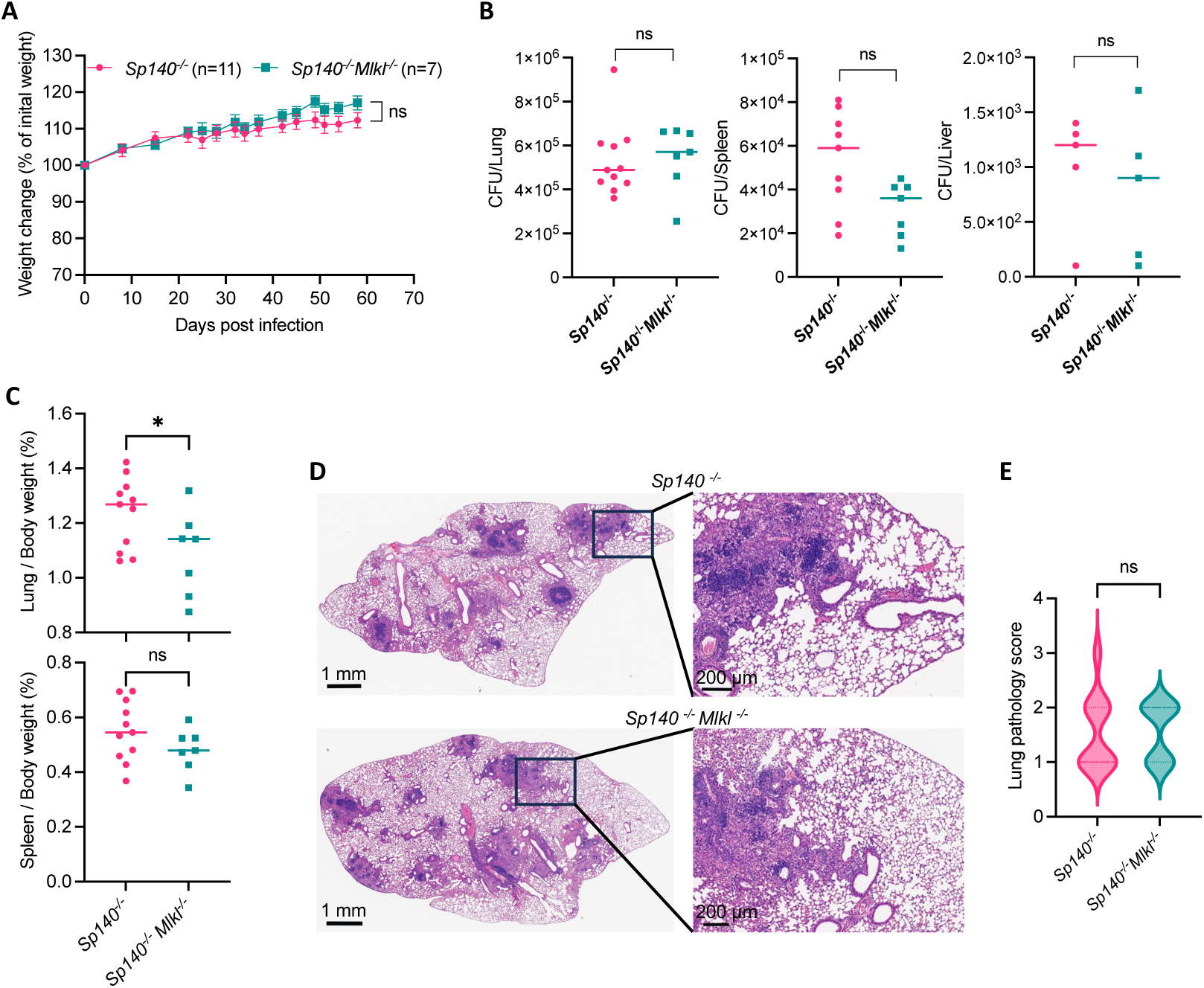
Medium dose of Mtb infection in *Sp140*^−/−^ mice results in slow TB progression, related to Figure 5. . *Sp140*^−/−^ and *Sp140*^−/−^*Mlkl*^−/−^ mice were infected with 400-500 CFU of Mtb on day 0. For (B-F), mice were euthanized on day 58. (A) Weight changes of Mtb-infected mice. (B) Mtb burdens in lungs (left), spleens (middle) and livers (right) of mice. (C) The ratio of lung (up) and spleen (down) weight to whole body weight in Mtb-infected mice. (D) Representative HE staining of lung sections of Mtb-infected mice. (E) Pathology score of lung sections of Mtb-infected mice. n=11 for *Sp140*^−/^ mice and n=7 for *Sp140*^−/−^*Mlkl*^−/−^ mice. Data are means ± SEM (A-C and E). ns, not significant; *, p < 0.05.(two-tailed unpaired Student’s *t* test).

